# Filament Rigidity and Connectivity Tune the Deformation Modes of Active Biopolymer Networks

**DOI:** 10.1101/141796

**Authors:** Samantha Stam, Simon L. Freedman, Shiladitya Banerjee, Kimberly L. Weirich, Aaron R. Dinner, Margaret L. Gardel

## Abstract

Molecular motors embedded within collections of actin and microtubule filaments underlie the dynamic behaviors of cytoskeletal assemblies. Understanding the physics of such motor-filament materials is critical to developing a physical model of the cytoskeleton and the design of biomimetic active materials. Here, we demonstrate through experiments and simulations that the rigidity and connectivity of filaments in active biopolymer networks regulates the anisotropy and the length scale of the underlying deformations, yielding materials with varying contractility. Semi-flexible filaments that can be compressed and bent by motor stresses undergo deformations that are predominantly biaxial. By contrast, rigid filament bundles contract via actomyosin sliding deformations that are predominantly uniaxial. Networks dominated by filament buckling are robustly contractile under a wide range of connectivities, while networks dominated by actomyosin sliding can be tuned from contractile to extensile through reduced connectivity via cross-linking. These results identify physical parameters that control the forces generated within motor-filament arrays, and provide insight into the self-organization and mechanics of cytoskeletal assemblies.

## INTRODUCTION

Assemblies of semi-flexible filaments and molecular motors are active materials (1) that drive many physiological processes such as muscle contraction (2), cytokinesis (3), cytoplasmic transport (4), and chromosome segregation (5). To actuate these processes, the nanometer-scale displacements of motors and local deformation and sliding of filaments must give rise to coordinated mesoscale deformations of such active materials. These mesoscale dynamics result in the transmission of cellular-scale forces with different directions (e.g., contractile or extensile) and shapes (e.g., isotropic or anisotropic) which, in turn, result in shape changes at cellular and ultimately tissue length scales. Characterizing deformations in active networks of different molecular compositions is a much needed first step toward understanding complex force transmission and shape changes observed in cells and tissues.

Understanding how assemblies of filaments and motors produce a net contractile or extensile force has been extensively explored theoretically (6–12). Experimentally, *in vitro* networks constructed from actin filaments and myosin II motors are robustly contractile (13–16). By contrast, systems of microtubules and molecular motors are either extensile (6, 7, 17, 18) or contractile (19, 20). One difference between these two active materials is that microtubules are significantly more rigid than actin. Recent work has shown that contractile stress can be generated via motor stress-induced filament buckling (11, 12, 15), indicating an important role for filament rigidity. Alternative microscopic mechanisms to generate extensile or contractile stress by motor-mediated sliding of rigid filaments have also been proposed (6–10). The network-scale consequences of these different force-generating mechanisms have not been explored.

Deformations within active matter can be characterized beyond whether they are contractile or extensile. For example, network-scale force transmission is known to be affected by network connectivity, which regulates the length scale of contraction (13, 14, 21–23). Moreover, recent data suggests that disordered actomyosin networks contract isotropically (24, 25). *In vivo*, anisotropic contraction dominates in cell division and muscle contraction (26). Understanding how to control the prevalence of isotropic versus anisotropic deformations will further our understanding of how these contractile deformations are regulated *in vivo*.

Here, we directly vary the stiffness and connectivity of filaments within an *in vitro* biopolymer network through cross-linking and investigate the effects on network deformation. Through quantitative analysis of experimental data, we determine that these mechanical properties affect the anisotropy and contractility of deformations caused by the motor protein myosin II. Networks composed of semi-flexible filaments that can be buckled by motor stresses exhibit robust biaxial contraction. Increasing the filament rigidity results in uniaxial deformations, the direction of which is regulated by cross-linker density. Extensile deformations are generated at low cross-linker density and contractile deformations occur at high cross-linker density. Using agent-based simulations, we identify the microscopic deformation modes underlying these observations and find that forces are transmitted uniaxially by rigid filaments that slide and do not buckle. Together, our results indicate how motor-filament interactions can generate forces that result in either extensile or contractile deformations, which vary in shape depending on the filament rigidity and connectivity. From our experimental and simulation data, we propose a phase space of active matter constructed from motors and filaments.

## RESULTS

### Networks of cross-linked rigid bundles are contractile with a short correlation length

To investigate the role of filament rigidity in active motor-filament networks, we construct a quasi-two-dimensional (quasi-2D) layer of actin *in vitro* by polymerization of 1 μM monomeric actin in the presence of a depletion agent to crowd actin filaments near a passivated surface (Fig. 1A) (15, 17). To increase filament rigidity, we add 0.1 μM of the actin cross-linker fascin, which constructs bundles of ~8 ± 7 actin filaments (Supplemental Fig. S1). Actin filaments are polar, and their barbed ends are uniformly directed within fascin bundles (27). Fascin bundles are thus polar like single actin filaments but are much more rigid (Fig. 1B): the persistence length of bundles is estimated to be ~250 μm (28), over 10 times that of single actin filaments (29). To connect rigid bundles into networks, we add a small concentration (0.002 μM) of a second crosslinker, filamin. Filamin is a large (200 nm) and flexible cross-linker that binds overlapping bundles with varying orientations into a quasi-2D network (30, 31).

**Figure 1:**
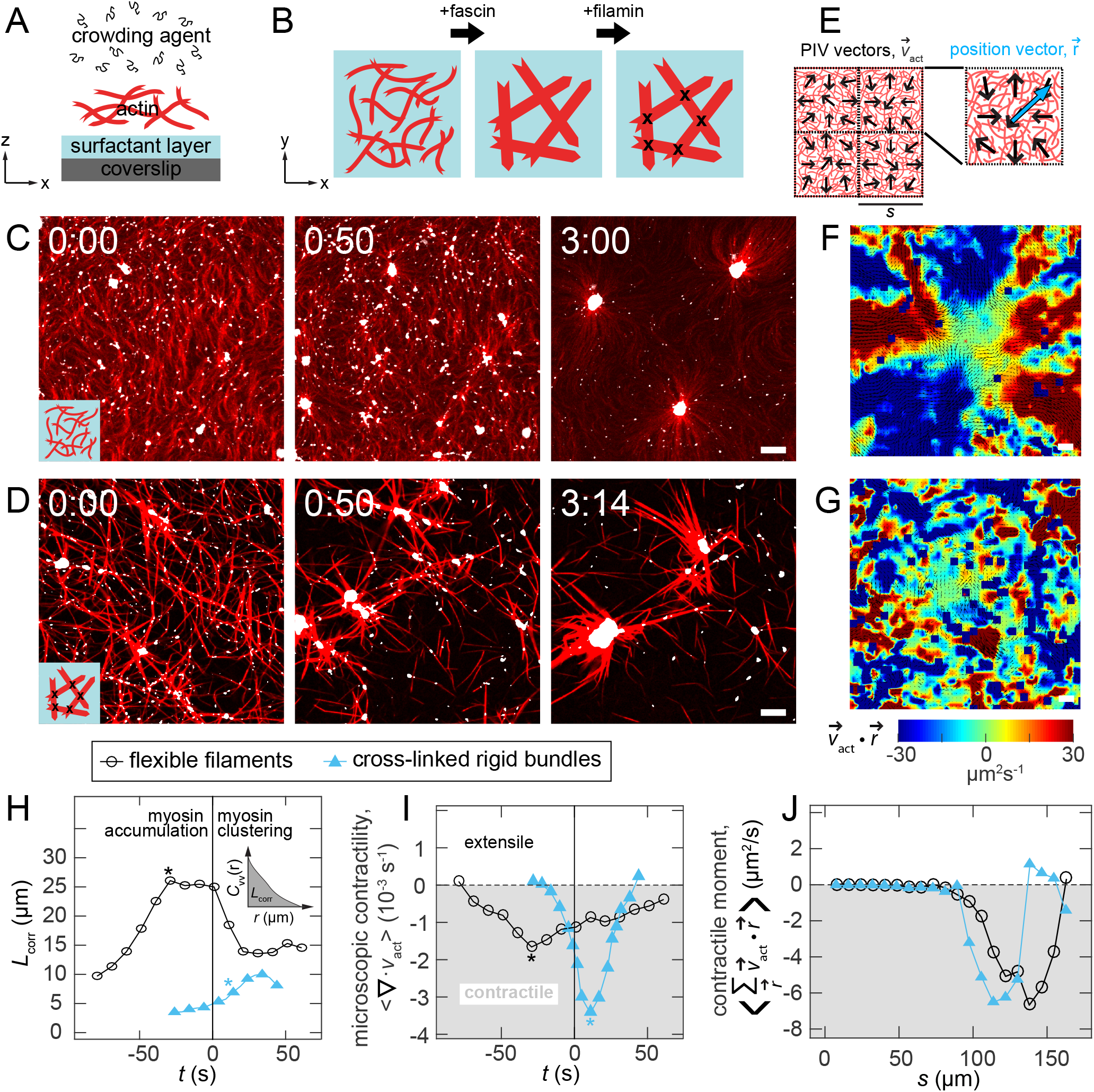
Networks of rigid bundles are contractile with a short correlation length. (A) Schematic of experimental set up. Actin filaments are crowded to a surfactant-coated coverslip surface to make a dense quasi-2D layer. (B) Fascin is used to make rigid, unipolar actin bundles. Filamin is used to cross-link bundles. (C) Images of semi-flexible filaments (red) in the absence of fascin or filamin after the addition of myosin (white puncta). (D) Images of cross-linked rigid bundles formed by F-actin in the presence of fascin (1:10) and filamin (1:500) after myosin is added. (E) Particle Imaging Velocimetry (PIV) detects local motion of F-actin (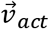, black arrows). Images are split into boxes of size *s*, and 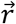 defines a vector from the center of a box to a PIV vector within the box. (F-G) Example spatial maps of the moment of the velocity field for images at −0:40 and 0:00 of panels C & D, respectively. Negative values of 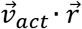 indicate contractile whereas positive values indicate extension. (H) The correlation length as a function of time for single filaments (open black circles) and cross-linked rigid bundles (closed blue triangles). Inset: Schematic indicating how correlation length is obtained from velocity-velocity correlation. (I) The divergence for both networks as a function of time. The asterisks in H+I indicate the time of minimal divergence, as indicated in (I). (J) The contractile moment as a function of length scale s for both samples. For (C)-(G), scale bars are 10 μm and time stamps are in the minutes:seconds format where 0:00 indicates the time of the maximal density of myosin puncta.

After assembling actin filaments or bundles, we add myosin II and monitor structural changes in the actin networks via fluorescence microscopy (Methods Section). Myosin II filaments (white spots) initially accumulate on the networks, and we define the time of the maximum density of myosin puncta as *t* = 0 s (Supplemental Fig. S2). Myosin drives changes in actin filament or bundle orientation, position, and shape that ultimately result in the formation of actomyosin asters comprised of polarity-sorted actin filaments oriented radially with large myosin foci at the center (Fig. 1C and 1D, Supplemental Fig. S2, Supplemental Movie S1 and S2).

To assess the network motion leading to aster formation, we calculate local displacement vectors of the actin network between frames using particle image velocimetry (PIV) (Fig. 1E, Methods). To visualize propagation of contractile or extensile motion, we calculate the moment of the velocity field, 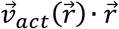, where 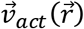 is the local actin PIV vector and 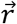 is the vector from the center of a square region to the location of the PIV vector (32) (Fig. 1E). Locations where the moment is positive indicate local expansion from the center of the field of view whereas negative values indicate local compression. During the early stages of network reorganization before aster formation, we find that spatial propagation of inwardly or outwardly directed motion is very different in networks of semi-flexible filaments and those of cross-linked rigid bundles (Figs. 1F and 1G). In networks of semi-flexible filaments, motion is highly spatially correlated, with large areas contracting toward the center of the square region in the vertical direction (blue, Fig. 1F) and material moving outward in the horizontal direction (red, Fig. 1F). In contrast, in the bundled network, motion is restricted to smaller, irregularly shaped contractile and extensile regions that are interspersed (Fig. 1G).

To characterize the length scale of the velocity field, we consider the velocity-velocity correlation function:

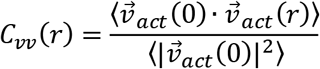

where *r* is the distance between two velocity vectors 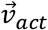. We define a characteristic correlation length, *L*_corr_, as the area under the curve of *C*_vv_(*r*) at a given time (inset, Fig. 1H). In both networks, *L*_corr_ initially increases as myosin forces accumulate in the network (Fig. 1H). Eventually, *L*_corr_ decreases as the networks break into clusters. Although *L*_corr_ has similar trends for both networks, its value is consistently less for the rigid bundle network than for the network of semi-flexible filaments. This is consistent with the spatial heterogeneity in the moment of the velocity field observed in the network of rigid bundles, as compared to that formed with semi-flexible filaments (Figs. 1F and 1G).

Next, we assess net contractility using two different measures. The divergence of 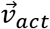, 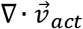, is a measure of contractility on the length scale set by the spacing of PIV vectors, in this case 2.4 μm (33). Negative values indicate local contraction while positive values indicate local expansion. For networks of semi-flexible filaments, the spatial average of 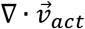 is negative (Fig 1I, open black circles), indicating net contraction, consistent with previous reports (33). The divergence reaches a maximally negative value as myosin accumulates on the network before separation of actin into clusters, at which point local extension between clusters balances contractility to produce 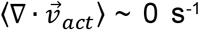. Similarly, the cross-linked rigid bundle network exhibits a negative divergence that returns to values near 0 s^−1^ after the onset of network coarsening at 0 s (Fig. 1I, filled blue triangles). Thus, the contractility is slightly enhanced in networks of rigid bundles as compared to those of semi-flexible filaments.

To characterize the length scale of contraction, we measure the contractile moment by summing 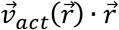 over non-overlapping square regions of varying side length *s* (Fig. 1J) (32). Negative values of the contractile moment indicate that contractile motion propagates across regions with this length scale (32). In both networks, 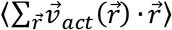 reaches a minimum for regions of length 100 μm < s < 150 μm. Thus, contraction in both materials can propagate over large length scales. However, the consistent picture that emerges is that the collective motions in the rigid networks occur over shorter length and time scales.

### Rigidity controls the anisotropy of contractile deformations

To explore the origin of differing spatial distribution of motion within these contractile networks, we sought to characterize the local deformations. We apply a method previously used to characterize the anisotropy of forces exerted by cells (32). We consider the tensor

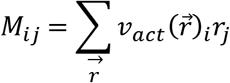

where *i* and *j* denote the in-plane spatial coordinates. By diagonalizing this tensor, we can extract the principle deformation axes. The magnitudes of the eigenvalues, *M*_max_ and *M*_min_, are the major and minor axes respectively of an ellipse characterizing the anisotropy of the deformation (Fig. 2A). A value of *M*_min_/*M*_max_ of 0 indicates a completely uniaxial deformation, while a value of *M*_min_/*M*_max_ = 1 indicates a completely biaxial deformation (Fig. 2A). For a given length scale (s = 20 μm), a distribution of *M*_min_/*M*_max_ from deformations across the field of view is obtained at each time point (Fig. 2B, Fig. 2C). In networks of semi-flexible filaments, the distribution is clearly weighted towards biaxial deformations (*M*_min_/*M*_max_ > 0.5) at all times during contraction (Fig. 2D). By contrast, in cross-linked rigid bundle networks, the distribution is highly weighted towards uniaxial deformations (*M*_min_/*M*_max_ < 0.5) at all times (Fig. 2E). We find that these characteristic differences in deformation anisotropy between rigid and semi-flexible networks persist across length scales varying from s = 6 μm up to 60 μm (Fig. S3).

**Figure 2:**
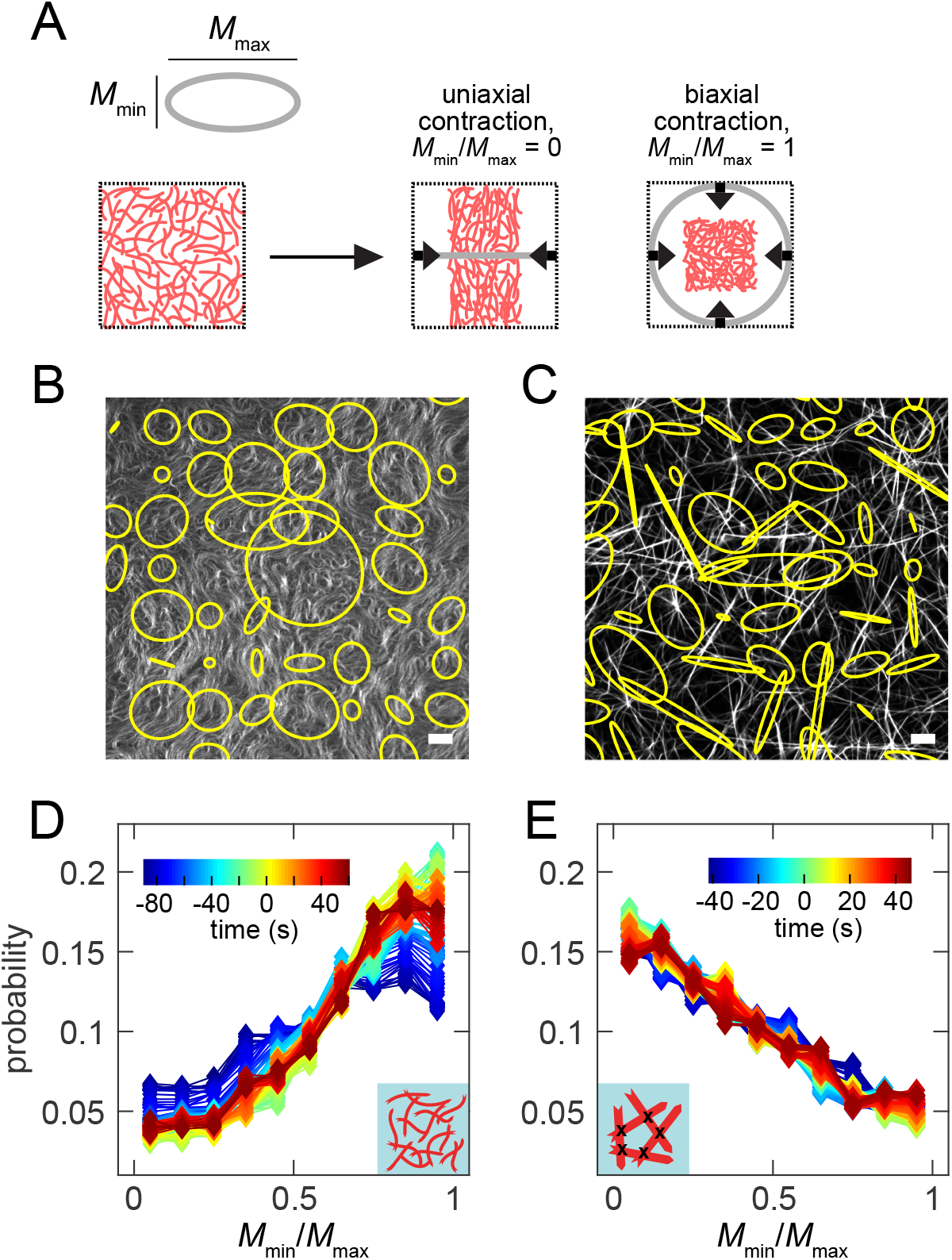
Deformations are highly biaxial and uniaxial in networks of semi-flexible filaments and rigid bundles respectively. (A) The eigenvalues of the flow dipole moment tensor, *M*_min_ and *M*_max_, are the axes of an ellipse that characterizes the deformation anisotropy, with uniaxial and biaxial contraction illustrated. (B) and (C) Images of deformation anisotropy in networks of semi-flexible filaments (B) and rigid bundles (C). (D) and (E) Distribution of *M*_min_/*M*_max_ at varying times (color scale) at s = 20 μm for semi-flexible filaments (D) and cross-linked rigid bundles (E).

To examine the effect of different deformations on correlated motion and contraction, we next consider the change in the fraction of predominately biaxial (*M*_min_/*M*_max_ > 0.5) or uniaxial (*M*_min_/*M*_max_ < 0.5) deformations and term these *P*_biaxial_(*s*) and *P*_uniaxial_(*s*) = 1 – *P*_biaxial_(*s*), respectively (Fig. S4). We compare these quantities to the correlation length, *L*_corr_, and the microscopic contractility as a function of time. Versions of these quantities that are rescaled to range from 0 to 1 are indicated by lower case letters, e.g., *p*_biaxial_(*s*) and *l*_corr_ (see Methods). For both rigidities, either *p*_biaxial_(*s*) or *p*_uniaxial_(*s*) is positively correlated with *l*_corr_ and is optimized for a given length scale *s* (Methods, Fig. S4). In networks of semi-flexible filaments, *p*_biaxial_ is positively correlated with *l*_corr_ during network contraction (Fig. 3A). In contrast, for the cross-linked rigid bundle networks, *p*_uniaxial_ is strongly positively correlated with *l*_corr_ (Fig. 3B and Fig. S4B). Interestingly, there is a time lag from the maximal contraction (minimum divergence) to the maximum *l*_corr_ and *p*_uniaxial_, which may arise from the higher sensitivity of the divergence measurement to biaxial contraction, as compared to uniaxial contraction.

**Figure 3:**
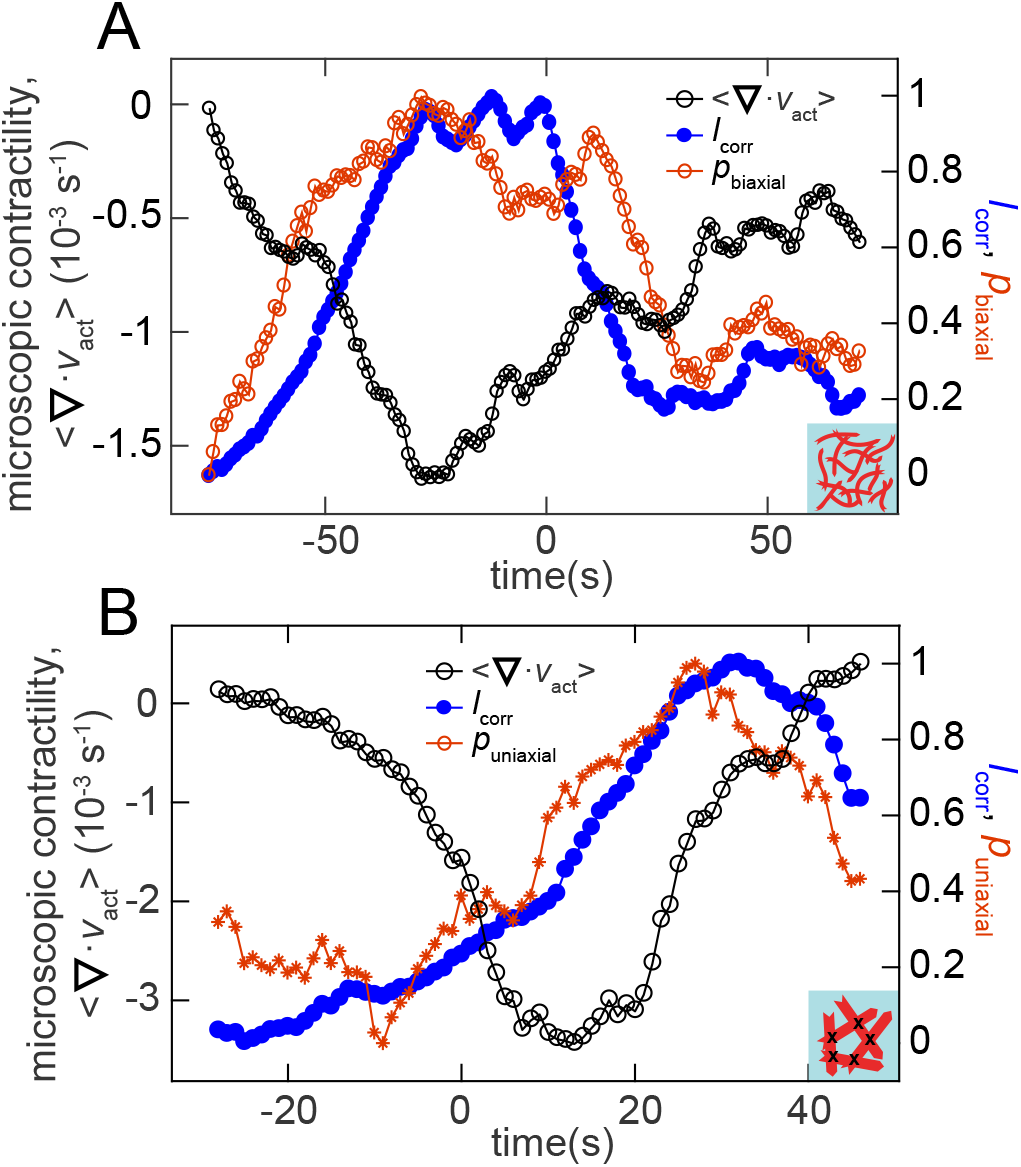
Activation of biaxial or uniaxial deformations in semi-flexible filament and rigid bundle networks respectively coincides with correlated motion and contractility. Plot of the divergence (open black cricles), correlation length and either biaxial probability (A) or uniaxial probability (B) as a function of time for single filaments (A) and cross-linked rigid bundles (B). The length scale chosen to calculate biaxial or uniaxial probability is determined to be the optimal one, as shown in Fig. S4 and is *s* = 25–30 μm in (A) and 55–60 μm in (B).

These data demonstrate that contractility can occur in networks composed of either semi-flexible or rigid filaments, consistent with previous reports of contractility in cross-linked biopolymer networks of varying composition (13–16). Our analysis reveals significant differences, however, in the mesoscale shape changes induced within the two networks, with compliant networks supporting biaxial contraction and rigid networks supporting uniaxial deformations.

Previously, we identified filament buckling as the microscopic mechanism underlying contractility in networks of semi-flexible actomyosin (15), an inherently biaxial deformation process. Further evidence for the buckling-based contractility is found in Supplemental Movie S3, where we observe that the shift to increasingly biaxial deformations in the distribution of *M*_min_/*M*_max_ in Supplemental Fig. S3A occurs concurrently with the development of visible buckling and bending in F-actin. This shift does not happen in the network of cross-linked rigid bundles (Supplemental Movie S4). The mechanism underlying contractility in networks of rigid bundles is presumably different because buckling is suppressed by increased filament rigidity for a constant motor stress.

### Uniaxial contraction arises from actomyosin sliding arrested by cross-linker accumulation

To elucidate the microscopic deformation modes underlying contraction in networks with varying filament rigidity, we use agent-based simulations (34). In brief, we model actin filaments as wormlike chains interacting with cross-linkers and motors represented as linear springs with two sites (heads) that can attach and detach to the filaments via a Monte Carlo procedure. When attached, motor heads walk toward filament barbed ends at a load-dependent speed. We use Langevin dynamics to evolve each structural component of the assembly in response to internal forces. When parameterized as detailed in (34), this model captures a variety of experimentally observed trends with reasonable quantitative accuracy. We implicitly model bundling, corresponding to experimental fascin-bundled actin, by varying the persistence length of the actin filament (*L_p_*) between 25 and 250 μm. We explicity model cross-linking, corresponding to the experimental cross-linker filamin, by a spring with rest length 0.15 μm. Myosin miniflaments are modeled similarly, as springs with rest length 0.5 μm, unloaded speed *v*_0_ = 1 μm/s, and stall force 10 pN.

We initially examine networks with filament rigidities similar to either actin filaments (persistence length, *L_p_* = 12.5*μ*m, Fig. 4A, Supplemental Movie S5) or fascin bundles (persistence length, *L*_p_ = 250 μm, Supplemental Movie S6) and equal cross-linker densities (*ρ_xl_* = 1 *μ*m^−2^, Fig. 4B). Consistent with experiments, we observe that motors (white rectangles) move actin filaments and rearrange the filaments into asters. Both of these networks show comparable extents of contraction (Fig. 4C). Performing simulations over a range of filament rigidities (2.5 μm – 250 μm) and cross-linker densities (0 − 1 *μ*m^−2^) reveals that microscopic contractility is generally more sensitive to changes in cross-linker density than filament rigidity (Fig. 4C).

**Figure 4:**
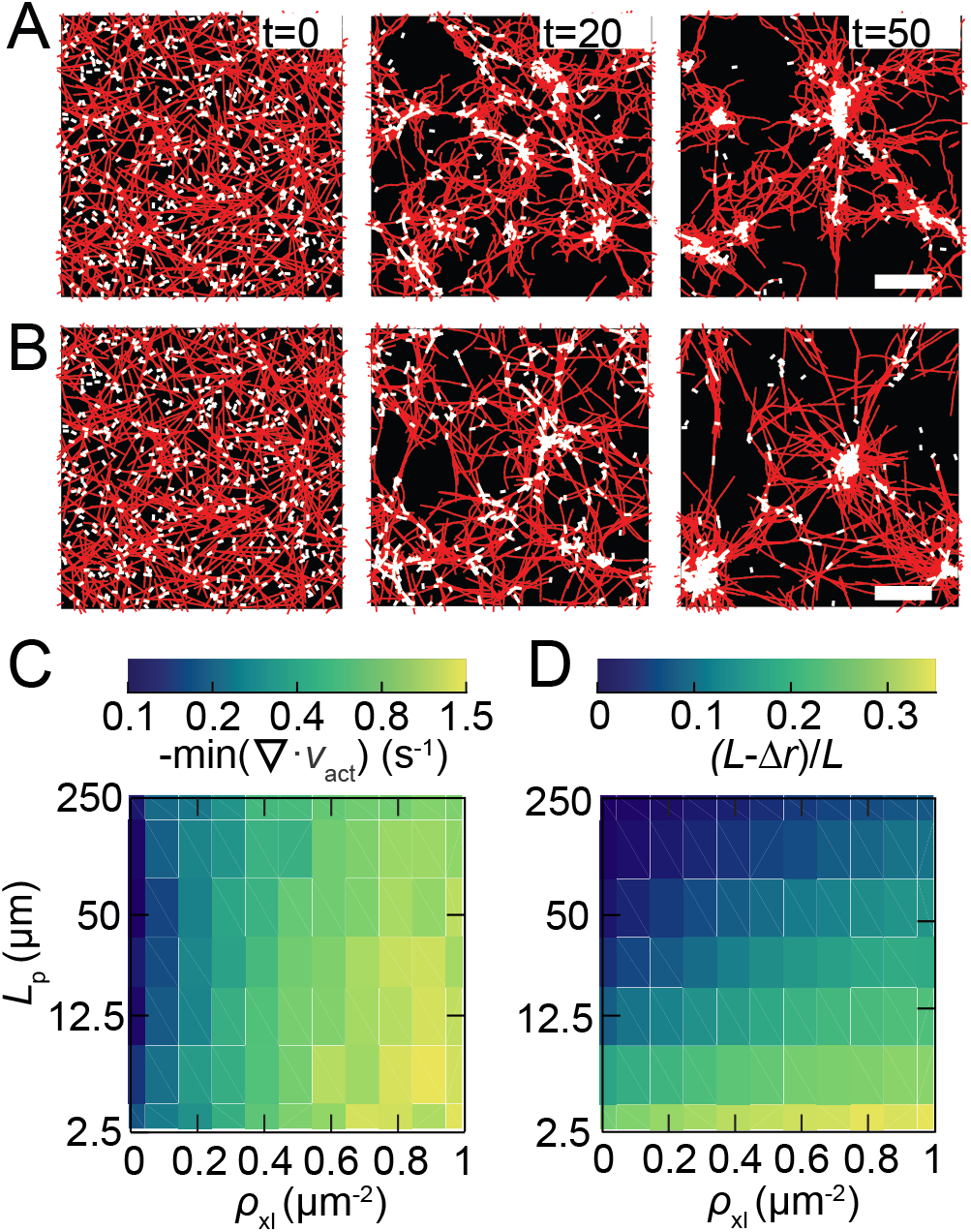
Simulations indicate cross-link dependent contractility over a wide range of filament stiffness. (A) and (B): Time series images in simulations for network with lower filament stiffness (*L*_p_ = 25 μm, (A)) and higher filament stiffness (*L*_p_ = 250 μm, (B)). Actin is shown in red and myosin is white. Scale bars are 10 *μ*m. (C): Microscopic contractility at varying filament stiffness and cross-link density. This is measured by the minimum of the spatially averaged divergence of the actin velocity field weighted by the local actin density in the first 25 s of simulation. (D) Filament compression during the first 25 s of simulation as a function of stiffness and cross-link density.

To explore the microscopic deformation modes underlying the regulation of contractility, we measure the filament deformation across these parameter values. One possible mechanism generating a net contractile deformation is filament buckling under local compressive forces (11, 12, 15). To quantify its extent, we measure the filament compression, *L − Δr*/*L* where *L* is the filament contour length and *Δr* is the end-to-end distance. This measure is zero when filaments are perfectly straight (*Δr* = *L*) and greater than zero if they are bent. The amount of compression is highest in the cross-linked networks comprised of flexible filaments (Fig. 4D). Filament compression decreases as the cross-linked density is lowered and approaches zero as the filament rigidity increases (Fig. 4D). Comparison of Figs. 4C and 4D shows that there is a sizable region of parameter space over which contractility occurs in the absence of filament compression. At our highest filament rigidities (*L*_p_ > 100 μm), contraction occurs with *L − Δr*/*L* less than 0.05 for *ρ_xl_* < 0.5 *μ*m^−2^. Lower filament rigidities require lower cross-linker densities to maintain small filament deformations.

An alternate microscopic mechanism of contractility that we expect to be pronounced at higher rigidities is myosin-driven actin sliding (6–9). Actin sliding drives local contraction when a motor connected to two antiparallel filaments is closer to their pointed ends, and local extension when it is closer to their barbed ends (Fig. 5A). In the absence of symmetry-breaking mechanisms, this would result in no net force propagation as extensile and contractile deformations would balance. However, when filaments overlap, there are more sites for cross-linkers to bind bivalently. This suppresses extensile motions that propagate force into the surrounding network (6). In the absence of cross-linkers, extensile motions can be favored by two mechanisms. First, for a uniform likelihood of myosin binding along the filament length, extensile antiparallel sliding will dominate (6, 7). Second, when the filaments reach the point of maximal overlap, they offer more available binding sites for motors to bivalently attach, which further increases extensile sliding (6, 7).

**Figure 5:**
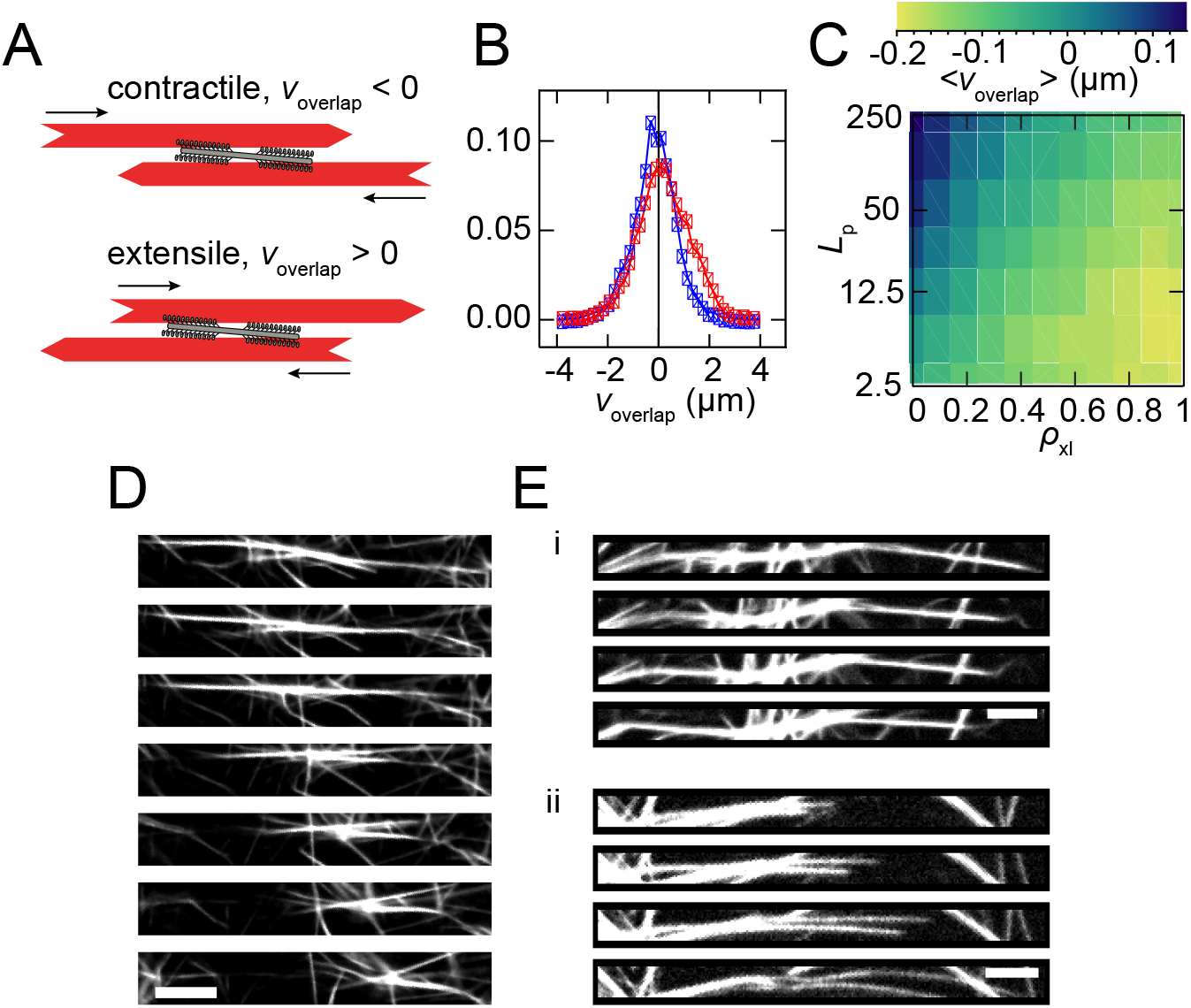
Uniaxial contractility is caused by arrested filament sliding. (A) A pair of antiparallel filaments are contractile if the myosin is near the pointed ends (top) and extensile is myosin is proximal to the barbed ends (bottom). (B) The distribution of *V*_overlap_ is shifted to more extensile values for rigid (*L*_p_ = 250 μm) filaments without cross-linking (red squares) compared to the same filaments with *ρ*_xl_ = 1 μm^−2^ (blue squares). This distribution is from the first 10 s of simulation. (C) Average of *V*_overlap_ over 25 s of simulation with varying filament rigidity and cross-link density. (D) In experiments with cross-linked rigid bundles, the bundles are observed to slide together and become arrested in the contracted state. The time delay between images from top to bottom is 1 s (E): In the absence of filamin, myosin drives both contractile (i) and extensile (ii) motions of rigid bundle pairs. The time delay between frames from top to bottom in both (i) and (ii) is 1 s Scale bars are 5 μm in (D) and (E).

We examine the probability distribution of relative sliding velocity, *V*_overlap_, in simulations of rigid (*L*_p_ = 250 μm) filaments both with (*ρ_xl_* = 1 *μ*m^−2^) and without cross-linkers. The distribution of overlap velocities shifts to negative values with the addition of cross-linkers (Fig. 5B). By examining the relative sliding velocity across all parameter values, we observe that the system is contractile (〈*ν_overlap_*〉 < 0) over most rigidities and cross-linker densities (Fig. 5C). However, at the lowest cross-linker densities and highest filament rigidities, we observe a regime where 〈*ν_overlap_*〉 > 0, indicating that extensile motions dominate.

To seek evidence for extensile sliding in our experiment, we examined pairs of bundles undergoing relative sliding. Indeed, in the presence of cross-links between bundles (1:500 filamin:actin) we observe bundle pairs sliding relative to each other, increasing the overlap, and then stopping (Fig. 5D). In a network without cross-links between rigid bundles, we see both relative motion between bundles that increases their overlap (Fig. 5E(i)) and relative motion that extends bundles further apart (Fig. 5E(ii)). The latter is similar to extensile motions observed in active liquid crystals of microtubules and kinesin (18), leading to the formation of asters (35, 36). Thus, our simulations and experiments of rigid filament suggests that cross-linker density can control the transition from contractile to extensile behaviors.

### Motors drive aster formation within rigid bundles without cross-links via uniaxial, extensile forces

To understand the consequences of the microscopic extensile deformations described above, we study the myosin-driven reorganization of rigid actin bundles that lack filamin cross-linkers but are sufficiently dense to have numerous overlaps such that myosin motors can slide and rearrange bundles to eventually form asters (Fig. 6A, Supplemental Movie S7). Asters are comprised of a dense myosin cluster with polarity sorted actin bundles emanating from the center, similar to those previously described (Figs. S2 and 6A). The spatial map of the moment of the velocity field reveals small contractile and extensile regions that are interspersed (Fig. 6B) and the velocity-velocity correlation length is short (Fig. S5). Consistent with simulations (Fig. 5C), the divergence of the velocity field indicates net extensile deformation (Fig. 6C), and the contractile moment is weakly positive at ~100 μm (Fig. 6D). The minimum divergence of the velocity field is weakly negative if the PIV vectors are calculated at sufficiently large time delays and length scales, but the divergence values are always less negative than in the other two networks (Supplemental Fig. S6). Consistent with motions dominated by actomyosin sliding, deformations are predominantly uniaxial (Figs. 6E, Fig. S3C, Supplemental Movie S8). Thus, actin sliding is responsible for short-range extensile, uniaxial deformations that drive local rearrangement of actin bundles into polarity-sorted asters.

**Figure 6:**
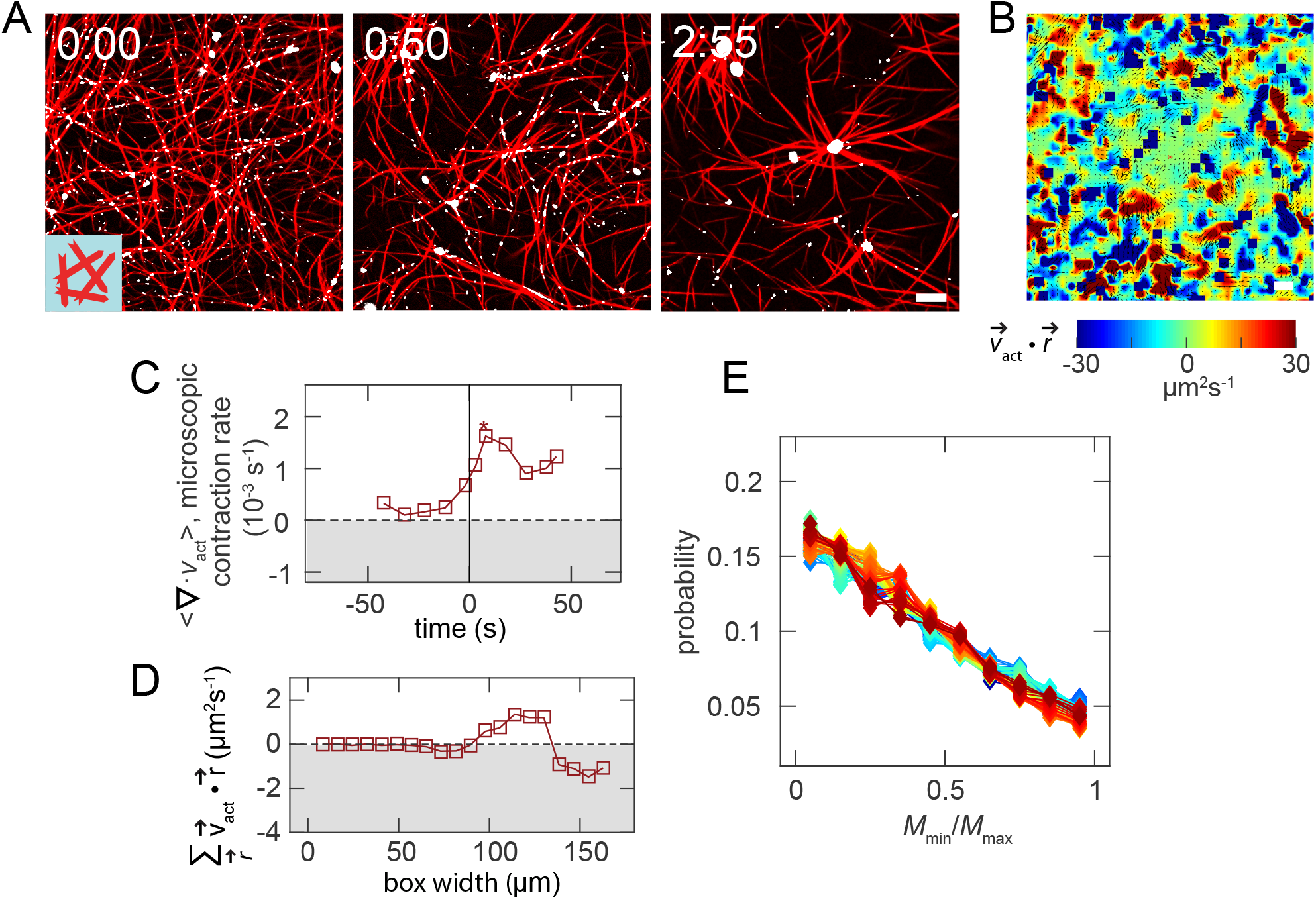
Myosin re-organizes rigid bundles via extensile lacking filamin cross-links via uniaxial forces. (A) Image sequence of fascin bundles without filamin. Actin is shown in red and myosin in white. (B): Values of 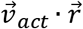 over a 150 μm x 150 μm square region. (C) The divergence of 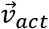 is non-contractile over the course of network rearrangement. (D) The contractile moment, 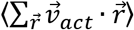, calculated over a 30 s interval after the maximum divergence in (C). (E) Distribution of *M*_min_/*M*_max_, s = 20 μm, indicates that deformations are predominantly uniaxial.

## DISCUSSION

Our results reveal three phases of deformation characterized by their anisotropy, length scale, and contractility. These can be controlled by modifying filament rigidity and connectivity in active biopolymer networks (Fig. 7). Moreover, we then demonstrate each phase is consistent with a unique microscopic deformation mode. In the presence of cross-linkers, we find that filament rigidity drives a transition between buckling-dominated and sliding-dominated contraction and, consequently, a transition between biaxial and uniaxial deformations. Such control over the shape of the deformations could be used to sculpt active materials both *in vitro* and *in vivo*. For rigid filaments, we find that increased cross-linking drives a transition from extensile to contractile deformation. While the role of cross-linking has been well described in terms of controlling force transmission (13, 14, 21–23), our work suggests that it also plays an important role in controlling the direction of the deformation, namely changing it from extensile to contractile. This result unifies previous observations of both extensile and contractile behaviors in active microtubule systems (6, 7, 17–20), suggesting that network connectivity is a significant factor in determining which behavior predominates. In future work, it will be interesting to explore the transitions between other microscopic deformation modes in active motor-filament systems and see how these are controlled by local structure or composition (e.g., filament orientation or polarity organization).

**Figure 7:**
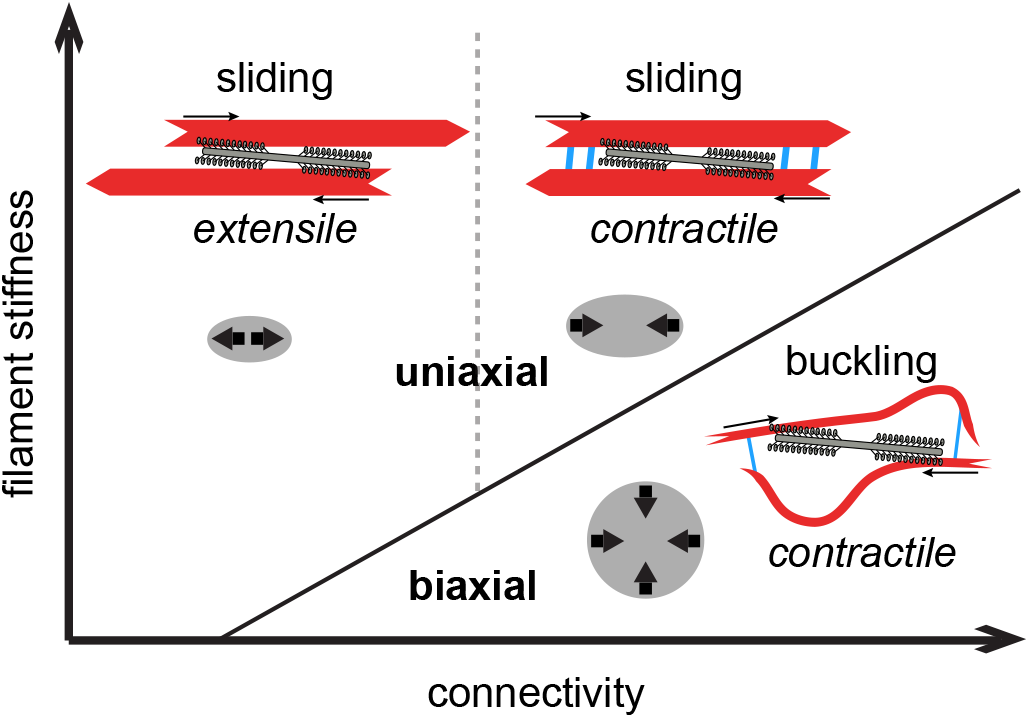
Uniaxial and biaxial deformations indicate differences in the mechanism of contractility and force propagation. Starting from the top left of the diagram, the three states we observe are extensile sliding, contractile sliding, and contractile buckling. The shape of the boundaries between these mechanisms are based on the simulation phase spaces in Fig. 4D and Fig. 5C. The mechanisms can be identified by the characteristic anisotropy of the transmitted forces, which is predominantly uniaxial for sliding and biaxial for buckling.

The myosin-driven remodeling of actin networks with varied connectivity and rigidity results in polarity-sorted asters of actin with high myosin densities at their centers, consistent with previous experiments reporting cluster formation (13–16, 19–21, 35–40). Many of these studies have equated cluster formation with contraction, and associated theoretical models have assumed that motors produce contractile force dipoles (41). However, our analysis shows that the microscopic driving forces and deformation modes to construct asters can include both isotropic and anisotropic contractility as well as anisotropic extension. While these generally result in differences in the actin distribution in the final structure (Supplemental Figure S2), our results show that different microscopic mechanics can result in similar final organizations. This underscores the importance of characterizing the dynamical rearrangements during active processes rather than relying on final structures alone to elucidate physical mechanisms.

Our work has implications for assessing and understanding the underlying physical mechanisms of force propagation in a variety of active biopolymer systems. Motor-filament arrays are a common motif in the actin and microtubule cytoskeletons during processes, including cell migration, cell division, intracellular transport, and formation of the mitotic spindle. Beyond the cytoskeleton, intranuclear molecular motors can drive correlated motion of chromatin (42), and forces produced by whole bacterial or mammalian cells can drive motions such as biofilm contraction or growth (43, 44) or alignment and organization of filamentous extracellular matrices (45–47). The physical properties of deformations that occur during these processes and the mechanisms at the level of biopolymer deformation or translocation have not been explored. Investigations of this nature will reveal which features of active matter dynamics are fundamental across these highly diverse systems and which features are regulated by particular biopolymer and motor network properties.

## Acknowledgements

This research was supported by the University of Chicago Materials Research Science and Engineering Center (National Science Foundation Division of Materials Research Grant 1420709). S.L.F. acknowledges support from the DoD through the NDSEG program. M.L.G. acknowledges support from National Science Foundation Molecular Cellular Biosciences Grant 1344203. S.S. acknowledges support from NIH NIBIB training grant T32EB009412. We thank Alyssa Harker and Dave Kovar for assistance with purifying fascin.

